# Gut microbiota plasticity is correlated with sustained weight loss on a low-carb or low-fat dietary intervention

**DOI:** 10.1101/580217

**Authors:** Jessica A Grembi, Lan H Nguyen, Thomas D Haggerty, Christopher D Gardner, Susan P Holmes, Julie Parsonnet

## Abstract

**Background:** Obesity is a complex global health challenge. Although both low-carbohydrate (low-carb) and low-fat diets can lead to weight loss, there is typically substantial variability in weight and related outcomes for both diet approaches among obese but otherwise healthy adults. Elucidating individual characteristics that might contribute to sustained weight loss is critical to developing effective dietary intervention strategies. We hypothesized that structural differences in the gut microbiota explained some portion of the weight loss variability among people randomized to either a low-carb or low-fat diet, possibly related to its effects on dietary compliance.

**Results:** Our study included two staggered cohorts of obese adults enrolled in the Diet Intervention Examining The Factors Interacting with Treatment Success (DIETFITS) study - a randomized clinical trial of either a low-fat or low-carb diet. In the discovery cohort (n=66), 161 pre-diet fecal samples were sequenced in addition to 157 samples collected after 10-weeks of dietary intervention. In the validation cohort (n = 56), 106 pre-diet fecal samples were sequenced. Pre-diet taxonomic features, such as the Prevotella/Bacteroides ratio, correlated to weight loss in the discovery cohort were not confirmed in the validation cohort. The most robust finding in the discovery cohort indicated that gut microbiota plasticity was linked to 12-month weight loss in a diet-dependent manner; subjects with higher sustained weight loss on a low-fat diet had higher pre-diet daily plasticity, whereas those most successful on the low-carb diet had greater microbiota plasticity over 10 weeks of dietary intervention. Unfortunately, because sample frequency and timing was quite different in the validation cohort, the relationship between plasticity and weight loss could not be studied in this group.

**Conclusions:** These findings suggest the potential importance of gut microbiota plasticity in sustained weight loss. We highlight the importance of evaluating kinetic trends and in assessing reproducibility in studies of the gut microbiota.

## Background

The global obesity pandemic has claimed one in three American adults and prevalences continue to rise in many other countries as well [46]. Obesity comorbidities (e.g. cardiovascular diseases, cancer, diabetes, and other chronic conditions) cost the US 8.65 billion/yr from loss of productivity due to absenteeism alone [58, 65, 53, 55, 14]. The personal, social and economic costs provide an urgent need for consistently effective, and possibly more personalized, weight-reduction therapies.

Different dietary interventions, such as low-carbohydrate (low-carb) and low-fat diets, can lead to weight loss, but not always; there remains substantial variability in diet success outcomes among obese, but otherwise healthy, adults [37]. Additionally, adherence to dietary intervention strategies has remained a major challenge, despite the clear dose-response relationship with weight loss for individuals on both low-carb and low-fat diets [17, 1]. Thus, practitioners have begun to look for individual characteristics (e.g. physiological attributes, cultural or lifestyle characteristics, food preferences, etc.) that could influence an individual’s sustained weight loss, possibly by improving adherence to a specific dietary regime.

Gut microbiota are highly individualized and intricately involved with the quantity and quality of nutrients extracted from our diets, with direct implications for obesity [19, 62, 26, 41, 60, 59]. Microbial metabolites and proteins are known to communicate with the host to influence appetite control [33, 11, 22, 49]. For example, proteins from gut *E. coli* modulate appetite by interacting with antigens involved in host satiety signaling [6]. Byproducts of microbial fermentation (e.g. butyrate and propionate) also stimulate gut hormones that reduce food intake [42, 50]. Studies assessing inter-individual variability in gut microbiota alterations in response to dietary interventions are limited by short-term (<12 weeks) dietary modifications or measurement of metabolic outcomes (e.g. plasma glucose, triglycerides, insulin, cholesterol) rather than weight loss [28]. The one published study that directly investigated the relationship between baseline microbiota structure and long-term weight loss found that a high fecal *Prevotella/Bacteroides* (P/B) ratio (> 0.01, n = 15) at baseline resulted in a mean of 1.31kg more weight loss at 6 months than a low P/B ratio (P/B < 0.01, n = 21) [32]. No studies have investigated the impact of microbiota on adherence to a dietary intervention.

The DIETFITS study [57, 24] was a randomized trial of 609 adults designed to elucidate predisposing individual characteristics–genotype, insulin-glucose dynamics, physiological and psychosocial attributes-that contribute to successful 12-month weight loss on *ad libitum* diets designed to be lower in carbs or fat. Within a population subset of DIETFITS, we explored whether attributes of the gut microbiota predisposed individuals to successful 12-month weight loss and how that might have been mediated by dietary adherence. Identifying pre-diet features of the gut microbiota that can predict adherence and/or success on a specific diet might permit personalization of dietary intervention strategies to maximize weight loss.

## Results

### Subject demographics and sequencing statistics

We recruited subjects from two cohorts of obese adults enrolled in the DIETFITS randomized trial of low-carb and low-fat diets. The cohorts were enrolled approximately six months apart, allowing one to be used for discovery and the second for validation. From each of these cohorts, individuals who a) provided fecal samples prior to initiating the intervention that passed quality filtering (>10,000 high-quality 16S rDNA sequences) and b) completed the one-year intervention were included in our study.

The discovery cohort included 66 subjects, of whom 32 (22 female) were randomized to the low-carb diet and 34 (17 female) to the low-fat diet. These 66 subjects provided fecal samples on three consecutive days prior to starting the diet plus three additional daily samples 10 weeks after diet initiation. A sequencing depth of 73, 659 ± 33, 380 reads per sample was obtained from 318 fecal samples (161 pre-diet; 157 at 10 weeks). Subject characteristics and dietary information can be found in Table ??. Subjects on the low-carb diet restricted carbs to 22.6 ± 10.3% of their daily kilocalories (kcals) and lost 8.4 ± 7.7% of their starting weight after 12 months of dietary intervention whereas those on the low-fat diet restricted fats to 25.3 ± 5.7% of daily kcals and lost 6.3 ± 7.7% of their starting weight. Previous definitions of long-term weight loss success [64, 36] were used to categorize subjects based on the percentage of baseline weight lost at 12 months: 20 were unsuccessful (US: < 3% weight loss), 25 were moderately successful (MS: 3 – 10% weight loss), and 21 were very successful (VS: > 10% weight loss).

**Table 1:** Subject characteristics and dietary intake for discovery and validation cohorts.

| width=1 |  |  |  |  |
| --- | --- | --- | --- | --- |
|  | Low-carb |  | Low-fat |  |
|  | Discovery<br>(n = 32) | Validation<br>(n = 31) | Discovery<br>(n = 34) | Validation<br>(n = 25) |
| Sex, n (%) |  |  |  |  |
| Female | 22 (68.8) | 25 (80.6) | 17 (50) | 19 (76) |
| Male | 10 (31.2) | 6 (19.4) | 17 (50) | 6 (24) |
| Age, yr (SD) | 43.1 (6) | 42.5 (6) | 40.5 (6.9) | 39.6 (6) |
| Race/ethnicity, n (%) |  |  |  |  |
| White | 25 (78.1) | 23 (74.2) | 25 (73.5) | 16 (64) |
| Hispanic | 4 (12.5) | 5 (16.1) | 7 (20.6) | 4 (16) |
| Asian | 3 (9.4) | 2 (6.5) | 2 (5.9) | 3 (12) |
| Other | 0 (0) | 1 (3.2) | 0 (0) | 2 (8) |
| Baseline weight, kg (SD) | 92.9 (16.8) | 90 (13.9) | 96.1 (12.4) | 92.6 (12.7) |
| Body mass index, kg/m<sup>2</sup> (SD) | 33.4 (3.7) | 32.8 (3.5) | 33.2 (3.3) | 33.2 (3.5) |
| Weight loss<sup>1</sup>, % (SD) | 8.4 (7.7) | 4.8 (6.2) | 6.3 (7.7) | 5.4 (7.6) |
| Weight loss<sup>1</sup> success category, n (%) |  |  |  |  |
| US | 9 (28.1) | 14 (45.2) | 11 (32.4) | 10 (40) |
| MS | 10 (31.2) | 11 (35.5) | 15 (44.1) | 7 (28) |
| VS | 13 (40.6) | 6 (19.4) | 8 (23.5) | 8 (32) |
| **Dietary Intake** |  |  |  |  |
| Fat, % kcal |  |  |  |  |
| Baseline | 38.9 (6.1) | 37.3 (6.4) | 36.8 (6.5) | 36.1 (5.2) |
| 3 months | 57 (9.1) | 48.3 (8.7) | 21.4 (7.2) | 24.9 (7.5) |
| 6 months | 53.6 (9.1) | 44.9 (7.5) | 25.6 (7.6) | 27.8 (9.1) |
| 12 months | 47.5 (10) | 43.2 (8.4) | 28.6 (8.2) | 31.3 (7.9) |
| Carbohydrates, % kcal |  |  |  |  |
| Baseline | 44.6 (7.6) | 45.9 (6.6) | 46.4 (7) | 46.8 (6.5) |
| 3 months | 17.6 (11) | 27.7 (10.5) | 59 (9.1) | 54.4 (9.3) |
| 6 months | 22.1 (11.7) | 31.4 (10.4) | 55.6 (8) | 52.3 (9.6) |
| 12 months | 27.3 (12.5) | 35.4 (10.2) | 53.3 (8.1) | 48.1 (8.5) |
| Protein, % kcal |  |  |  |  |
| Baseline | 16.6 (3.4) | 16.8 (2.9) | 16.8 (3.4) | 17.1 (3.4) |
| 3 months | 25.4 (6) | 24 (6.1) | 19.5 (4.6) | 20.7 (5.4) |
| 6 months | 24.3 (5.2) | 23.6 (6.8) | 18.7 (4.2) | 19.9 (5) |
| 12 months | 25.2 (7.1) | 21.4 (4.9) | 18.1 (3.9) | 20.5 (4.5) |
| **Dietary compliance<sup>2</sup>** |  |  |  |  |
| Portion of diet from non-restricted foods<sup>3</sup>, % kcals (SD) | 77.4 (10.3) | 68.5 (9.1) | 74.7 (5.7) | 72.3 (7.4) |
| Reduction in restricted foods<sup>4</sup>, % kcals (SD) | 21.9 (10.5) | 14.4 (10.4) | 11.4 (7.3) | 8.4 (6.1) |

<sup>1</sup> Measured after 12 months of dietary intervention<sup>2</sup> Averaged over 3-, 6-, and 12-month dietary recall periods<sup>3</sup> Carbohydrate restriction for subjects on low-carb diet; fat restriction for subjects on low-fat diet<sup>4</sup> Decrease from baseline levels

The validation cohort was comprised of 56 subjects: 31 (25 female) on the low-carb and 25 (19 female) on the low-fat diet. Subject characteristics were comparable between the two cohorts (Table ??), except for the percent weight lost at 12 months on the low-carb diet, which was significantly lower in the validation cohort (4.8±6.2% compared to 8.4±7.7% in the discovery cohort, Welch’s t-test *p* = 0.045) and percent of carbs consumed (31.5 ± 9.1% compared to 22.6 ± 10.3% in discovery cohort, Welch’s t-test *p* = 0.0005). Subjects were classified to weight loss success groups, as described above, with the following distribution: 24 US, 18 MS, and 14 VS. From these “validation” subjects, fecal samples were collected only prior to the start of the dietary intervention at a median of 12 (IQR 7, 25) days apart. Samples meeting quality criteria (n = 106) had a mean sequencing depth of 70,041 ± 15, 664 reads per sample.

### Pre-diet gut microbial community composition does not predict 12-month weight loss success

Pre-diet gut microbial community composition varied among subjects, and samples collected from the same individual tended to cluster together in principal coordinates analysis (PCoA) shown in Fig. 1. In the discovery cohort, pre-diet microbiota composition did not cluster by 12-month weight-loss success category (PERMANOVA on Bray-Curtis dissimilarity low-carb: *p* = 0.51; low-fat: *p* = 0.81). The microbiota composition also did not cluster by age, gender, pre-diet weight, body mass index or dietary compliance in the PCoA maps. Similar results were found in the validation cohort (Fig. 1b).

**Figure 1:**
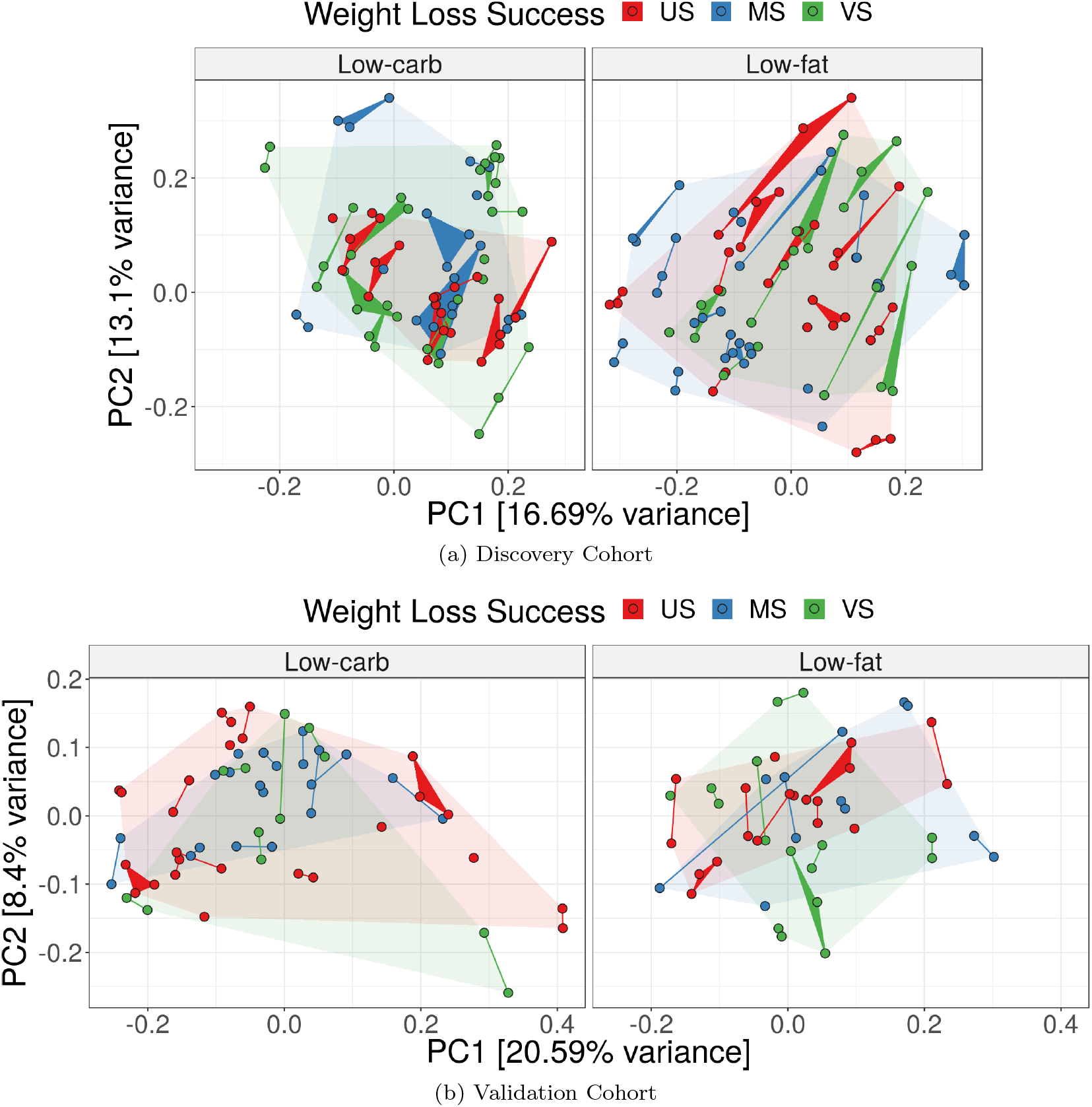
Pre-diet microbial community composition not correlated with weight-loss success. Pre-diet fecal microbiota collected from subjects in the discovery (a) and validation (b) cohorts prior to a low-carb (*left*) or low-fat (*right*) dietary intervention. Each point represents a single fecal sample and samples corresponding to the same subject are connected forming edges or triangles. Colors indicate 12-month weight-loss success: very successful (VS), moderately successful (MS), and unsuccessful (US). The faded background polygons show convex hulls for corresponding success categories. PCoA was computed with Bray-Curtis distance on inverse-hyperbolic-sine transformed counts.

### Higher gut microbiota plasticity predicted successful 12-month weight loss

In the discovery cohort, 85% of subjects (n = 26 for low-carb, n = 30 for low-fat) provided two or three fecal samples on consecutive days prior to the start of the intervention. These samples allowed us to quantify subjects’ *pre-diet daily microbiota plasticity*, i.e., the amount of daily variability in an individual’s microbiota composition, measured with *β*-diversity metrics. Pre-diet microbiota plasticity was significantly higher for VS subjects on the low-fat diet compared to US subjects (baseline BL v. BL plots in Fig. 2, Wilcoxon rank-sum test *p* = 0.033 for Bray-Curtis dissimilarity). This result was robust to the choice of distance metric (Fig S3) and was repeated on clusters of phylogenetically related ASVs (sharing roughly species- or genus-level sequence similarity; see Methods for details) with similar results (Fig S3b). There was no difference in pre-diet plasticity between weight-loss success groups for the low-carb diet. Consecutive daily samples were also collected after subjects had been on the dietary intervention for approximately 10 weeks; there was no difference in daily plasticity between weight-loss groups at 10 weeks for either diet (10wk v. 10wk plots in Fig. 2, Wilcoxon rank-sum test on Bray-Curtis dissimilarity *p* = 0.61 for low-carb, *p* = 0.54 for low-fat).

**Figure 2:**
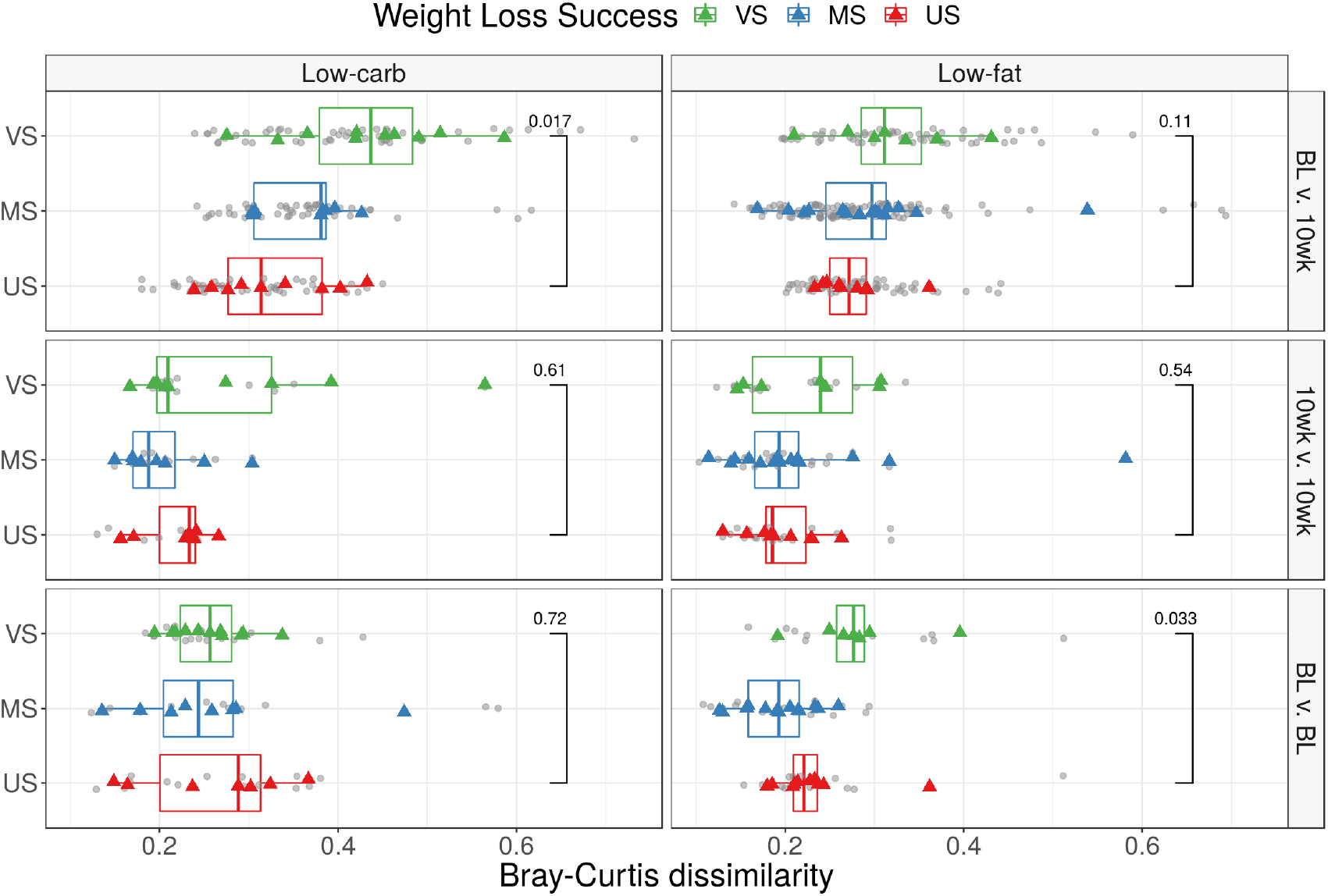
Gut microbiota plasticity over different periods. Pairwise *β*-diversity is shown between daily pre-diet samples (BL v. BL), between daily samples taken 10 weeks after initiation of the dietary intervention (10wk v. 10wk), and between BL and 10-week samples (BL v. 10wk). Bray-Curtis dissimilarities are shown for low-carb (left) and low-fat (right) diets. Grey points indicate computed pairwise dissimilarities between samples; colored points correspond to the average dissimilarity for each subject and are colored by weight-loss category: US – Unsuccessful, < 3% weight loss; MS – Moderately successful, 3 – 10% weight loss; VS – Very successful, > 10% weight loss. Results with other *β*-diversity metrics are shown in Figs. S3,S4

We also quantified *plasticity over ten weeks* in response to the dietary intervention, i.e., the variability between gut microbiota composition before the start of the dietary intervention and after 10 weeks of dieting. The plasticity over ten weeks was computed for subjects (n = 28 low-carb, n = 32 low-fat) who provided at least one pre-diet sample and another ten weeks after the start of the dietary intervention (77 ± 9 days apart for low-carb diet, 81 ± 14 days for low-fat). For each subject, the average pairwise *β*-diversity (between each pre-diet and 10-week sample) was calculated. On both diets VS subjects had higher plasticity between their baseline and 10-week fecal microbiota communities than US subjects (BL v. 10wk plots in Fig. 2; Wilcoxon rank-sum test on Bray-Curtis dissimilarity: low-carb *p* = 0.017; low-fat *p* = 0.11). Again, for both diets the observed trends were robust across multiple *β*-diversity metrics and clustered-ASV data (Fig. S4). Daily pre-diet plasticity was positively correlated with plasticity over ten weeks for subjects on the low-fat diet (Spearman’s rank correlation 0.37 for Bray-Curtis, *p* = 0.053). The magnitude of variation between the pre-diet and the 10-week period was higher than day-to-day plasticity at either the baseline or 10-week time point (Fig. 2, Wilcoxon rank-sum test *p* < 0.0001 for both low-carb and low-fat). Although intra-individual plasticity over ten weeks was significantly higher for VS compared to US subjects, inter-individual differences across subjects were still larger in magnitude on both diets (Fig. S5).

Within each cohort, pre-diet phylogenetic *α*-diversity was not different between US and VS subjects (Fig. S1) and was not correlated with dietary compliance on either diet (data not shown). However, for subjects in the discovery cohort, we found a negative correlation between pre-diet plasticity and average *α*-diversity (mean of all pre-diet samples) for subjects on the low-fat diet (Spearman’s rank correlation coefficient -0.42 for Bray-Curtis dissimilarity, p = 0.021). This suggests that individuals on the low-fat diet with low mean bacterial alpha-diversity tend to exhibit a more variable microbiota composition.

The magnitude of gut microbiota plasticity (using any *β*-diversity metric) was not statistically different between women and men at any time point: pre-diet (Wilcoxon rank-sum test on Bray-Curtis distance *p* = 0.86), at 10-week (*p* = 0.35), or between pre-diet and 10-week (*p* = 0.54). Subjects in the validation cohort did not collect pre-diet fecal samples on consecutive days (median 5 12 [IQR 7,25] days between samples) and fecal samples from 10-weeks into the intervention were not available, so we were unable to validate plasticity as a factor in weight loss success.

### Gut microbiota plasticity correlated with dietary compliance in a diet- and sex-dependent manner

Subjects were instructed to strive to restrict nonfiber carbohydrate or fat intake to 20g/d but to titrate over the first few weeks of the study to levels they perceived they could maintain for the rest of their lives. Each diet resulted in a reduced intake of the restricted component (Table ??) but also in the number of total calories consumed. Thus we calculated a proxy measure for adherence to a low-fat or low-carb diet, referred to here as *dietary compliance* and quantified as the reduction in restricted foods (measured as decrease in percentage of total daily kcals consumed from fats/carbs) between baseline (pre-diet) and during the dietary intervention (an average was taken over three unannounced 24-h dietary recalls at each time point conducted during the trial - baseline, 3, 6, and 7 12 months). We conducted sub-group analyses on dietary compliance for men and women, given observed sex-specific differences in correlations between reported dietary compliance and weight loss in the larger DIETFITS study population.

Women and men on the low-fat diet exhibited different correlations between dietary compliance and pre-diet daily plasticity (Fig. 3). Men with higher plasticity reduced their fat consumption more (had higher compliance) than men with lower plasticity (Spearman’s rank correlation using Bray-Curtis dissimilarity 0.55, p = 0.02), whereas women lacked a meaningful correlation between plasticity and dietary compliance (Spearman’s rank correlation -0.47, p = 0.1). For both women and men on the low-carb diet, no significant correlations were found between dietary compliance and daily pre-diet plasticity. These trends were consistent across several distance metrics (Fig. S6) and using ASV-clusters (data not shown).

**Figure 3:**
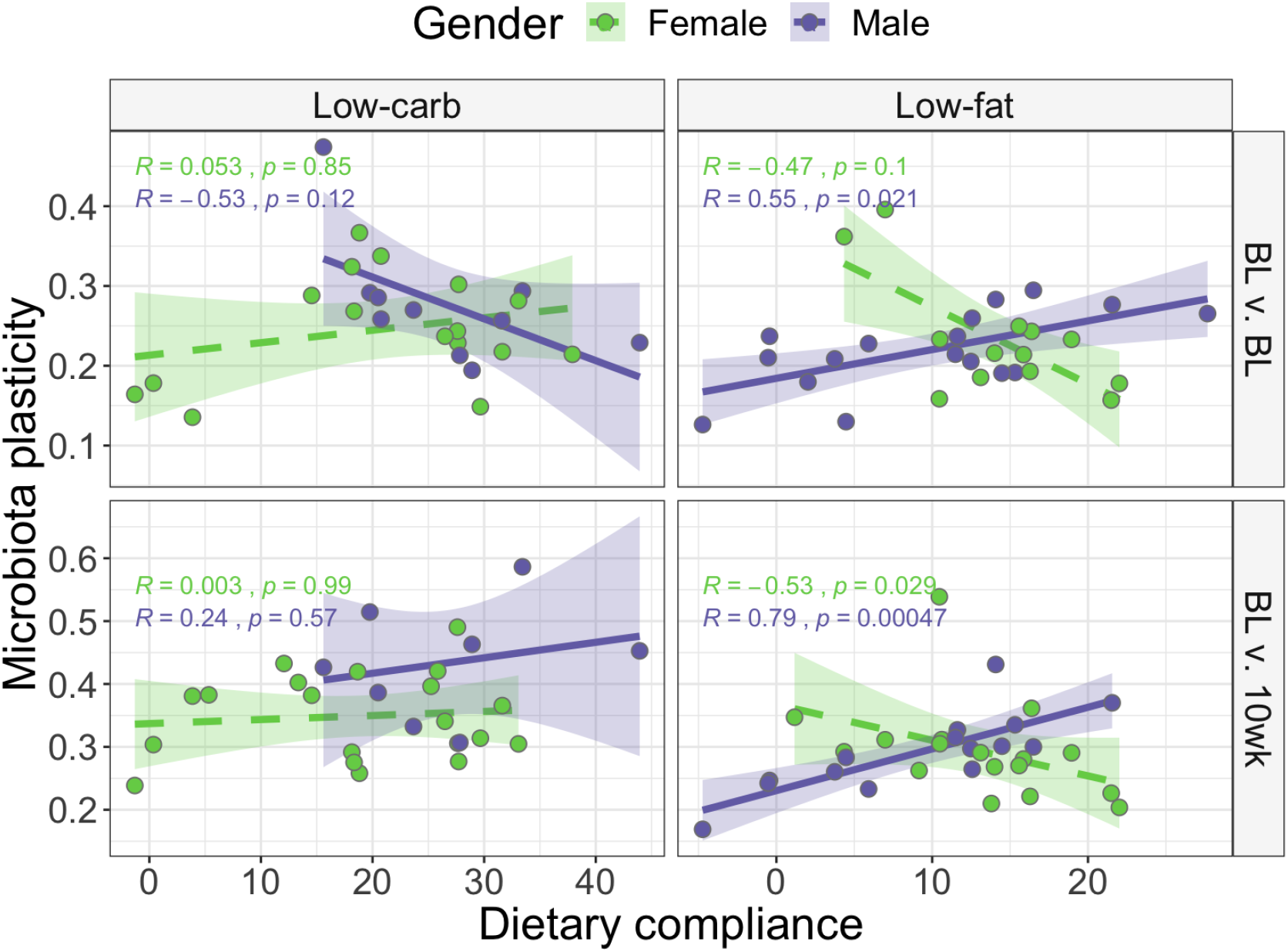
Gut microbiota plasticity correlated to dietary compliance in a sex- and diet-dependent manner. Spearman’s rank correlations between dietary compliance and plasticity (measured as Bray-Curtis dissimiliarity) between daily pre-diet samples (BL v. BL) and between pre-diet and 10-week samples (BL v. 10wk) are shown for low-carb (left) and low-fat (right) diets. Male (purple) and female (green) subjects show opposite correlations in many cases. Results with other *β*-diversity metrics are shown in Fig. S6.

We suspected that dietary compliance would affect diet-induced microbiota plasticity over ten weeks; more specifically, we expected the gut bacterial community composition of the more adherent subjects to shift more in response to their modified diet on the intervention). We observed highly significant and opposite correlations between men and women on the low-fat diet (see Fig. 3). This was not the case for subjects on the low-carb diet, where again no correlations were noted between diet-induced plasticity and compliance to the prescribed diet for both women and men. Trends were consistent across a variety of distance metrics (Fig. S6).

### Pre-diet Prevotella/Bacteroides ratio correlated with 12-month weight loss success on low-carb diet in discovery, but not validation, cohort

In the discovery cohort, subjects on the low-carb diet with VS weight-loss had significantly higher *Prevotella/Bacteroides* (P/B) ratio (median = 0.014) compared to US subjects (median = 0.0004; Wilcoxon rank-sum test *p* = 0.021); however, the same was not observed in the validation cohort (VS median P/B ratio = 0.0003; US median = 0.0009; *p* = 0.718; Fig. 4). There was no difference in P/B ratio between US and VS subjects randomized to the low-fat diet for either cohort (discovery: *p* = 0.54; validation: *p* = 0.46).

**Figure 4:**
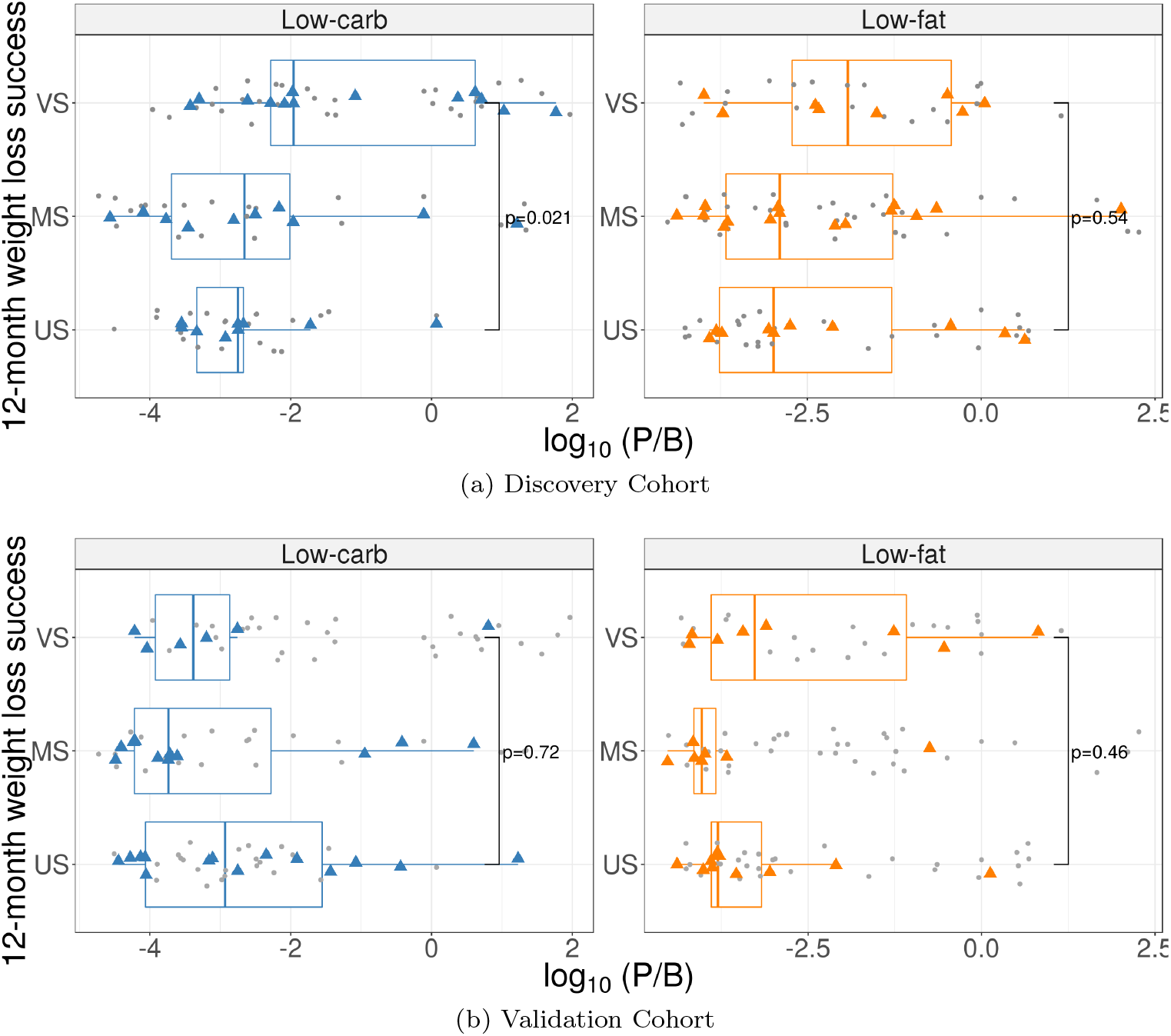
Differences in *Prevotella/Bacteroides* (P/B) ratio for subjects on a low-carb (left panels) or low-fat (right panels) diet. Grey points indicate P/B ratio for individual samples; colored points correspond to the average P/B ratio for each subject. Data from the discovery cohort (a) and the validation cohort (b) is displayed by subject’s weight-loss success category at 12 months after the start of the dietary intervention: US – Unsuccessful, < 3% weight loss; MS – Moderately successful, 3–10% weight loss; VS – Very successful, > 10% weight loss. P-values shown for Wilcoxon rank-sum test comparing US and VS groups.

### Differential abundance of sequence clusters not consistently predictive of 12-month weight loss success

To evaluate patterns of specific members of the pre-diet microbiota that might predict weight loss success, we tested for clusters of phylogenetically related ASVs (sharing roughly species- or genus-level sequence similarity; see Methods section for details on ASV-clustering) that differed in abundance between dieters who achieved VS 12-month weight loss compared to US. We compared within each diet separately, filtering out clusters that were not present in at least 10% of subjects from the focal diet. Using methods allowing for subject as a random effect (`limma` package, see Methods), in the discovery cohort we found one cluster containing 10 Ruminococcaceae ASVs (Cluster94) that was significantly more abundant in US compared to VS subjects on the low-carb diet (*p* = 0.0006) (see Table 2). In contrast, VS subjects had significantly higher abundances of a cluster containing 64 different Ruminococcaceae ASVs (Cluster65, (*p* = 0.023) and a cluster containing 2 *Enterorhabdus* ASVs (Cluster266, *p* = 0.025) compared to US subjects on the low-carb diet. The abundances do not display a proportional linear dose-dependent relationship with percentage weight loss for all subjects, however (Fig. 5), suggesting that these clusters are unlikely to be strong predictors of weight-loss success in a larger population. No clusters were identified as differentially abundant for the low-fat diet. Similar results were obtained when differential abundance testing was performed on the unclustered ASV counts (Fig. S7) and also when a linear model with a continuous predictor corresponding to the percentage weight loss (instead of categorical: VS vs US) was applied. ASVs and ASV-clusters identified as differentially abundant in the discovery cohort were not significant in the validation cohort (Table 2). In most cases, both the magnitude and direction of effect were discordant between cohorts.

**Figure 5:**
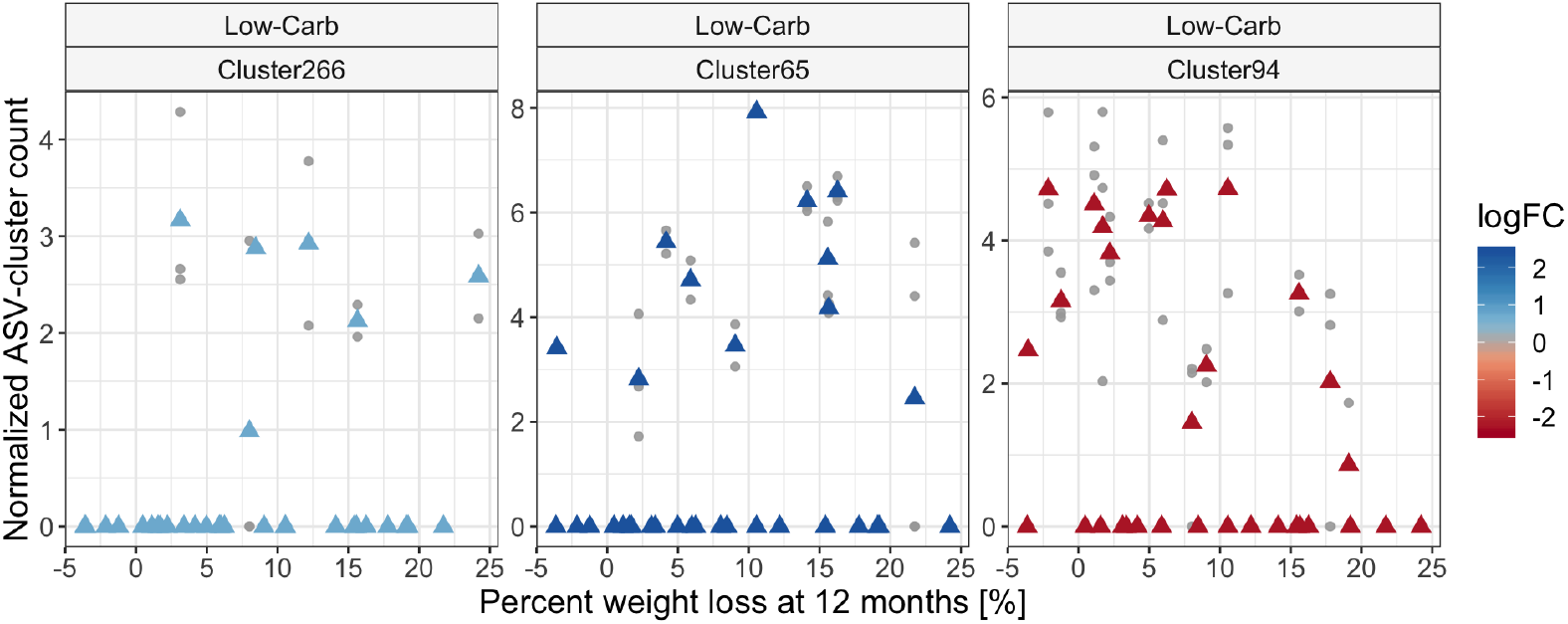
Taxa differentially abundant between weight loss success groups. ASV-clusters found differentially abundant when comparing subjects that were VS compared to US at 12-month weight loss on the low-carb diet. ASV-clusters were normalized and asinh-transformed for variance stabilization prior to analysis; the normalized, transformed values are shown on the y-axis. ASV-clusters have a median 96.8% sequence similarity (a taxonomic description can be found in Table 2). No taxa were found differentially abundant on the low-fat diet. Grey points represent individual samples and triangles represent the mean value for each subject.

**Table 2:**
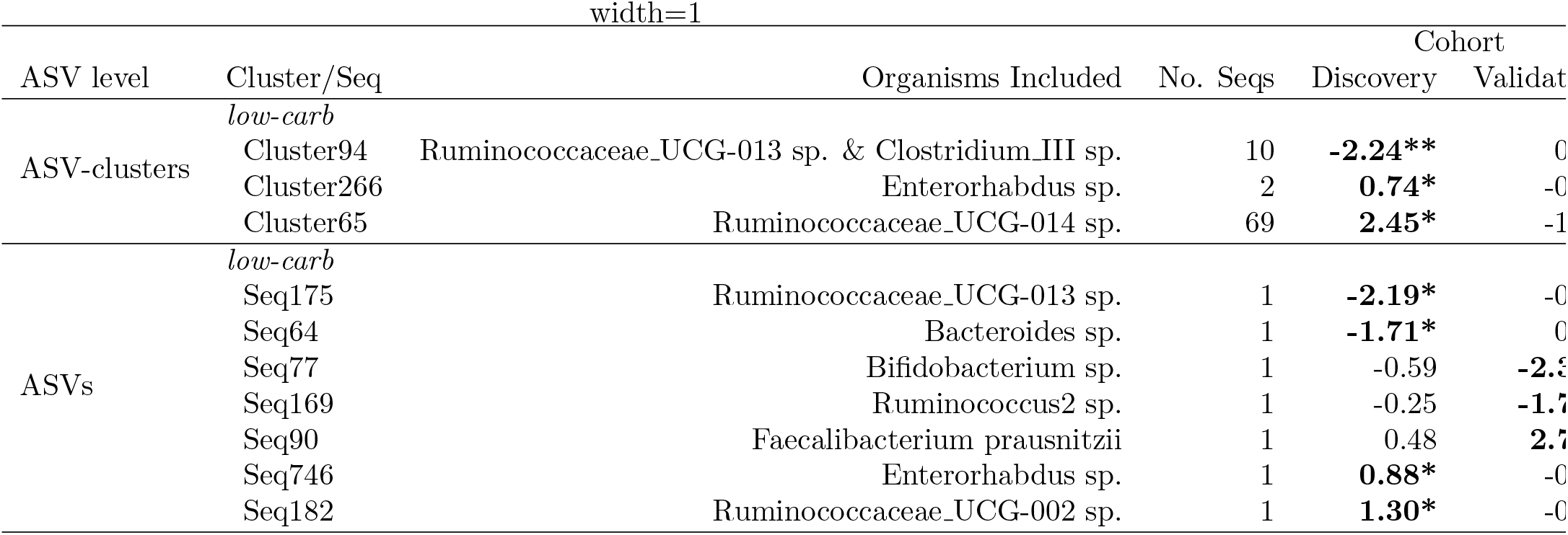
Log-fold changes of ASVs and ASV-clusters found differentially abundant when modeled against weight-loss success category at 12 months. Values represent log-fold changes between the US and VS groups. Bold values were significant for the cohort indicated. No clusters or ASVs were found differentially abundant in either cohort for the low-fat diet. * = *p* < 0.05, ** = *p* < 0.001

## Discussion

Our study suggests the potential importance of gut microbiota plasticity in sustained weight loss. The observed patterns in correlations between plasticity and our estimates of dietary compliance could imply sex- and diet-dependent mechanisms. In several earlier studies [15, 20, 68], individual temporal variability of the gut microbiota was seemingly eclipsed by larger-scale variability across body habitats, severe perturbations, or geographically distinct populations, which may have led some to underestimate the importance of plasticity within an individual. However, a study of weekly samples from 85 adults showed that temporal variability is a personalized feature [23]; with some individuals displaying consistently high or low compositional variability. Our results are the first to illustrate that this personalized feature of the microbiota might be relevant to weight loss.

Higher pre-diet daily plasticity in gut community composition was observed in subjects attaining higher 12-month weight loss, but only on the low-fat diet. This result was consistent across several *β*-diversity metrics indicating that microbiota was more plastic both in terms of membership and structure. Of note, no pre-diet microbes were differentially abundant for US compared to VS subjects on the low-fat diet, a result that might have been influenced by the increased plasticity found in the latter. In agreement with other studies, we saw a negative correlation with compositional variability and phylogenetic diversity (more diverse communities were less variable) [23, 16, 44]. The insurance hypothesis in ecological theory suggests that biologically diverse communities are more resilient as they contain a larger set of community traits/functions that enable them to adapt to changing environments and buffer the system against the loss of species [67]. In our case, we see that higher turnover of microbes might allow for a greater response to the dietary intervention, independent of diversity, possibly by allowing for the alteration of the microbial consortia into one less efficient at extracting nutrients.

As this was seen only for low-fat dieters, we hypothesized that increased plasticity might have facilitated responsiveness to the increased carbohydrates and fiber consumed on the low-fat diet [24], making the transition to the new diet easier for subjects in ways that aided adherence through appetite suppression [33, 39, 8, 25, 12, 3, 2]. Our data suggest this is true for men, as we saw positive correlations between plasticity and both weight loss and dietary compliance. Despite magnitudes comparable to men’s daily pre-diet plasticity, women on the low-fat diet displayed a negative trend between plasticity and dietary compliance. This suggests that the mechanisms underlying the relationship between plasticity and weight loss might be independent of dietary adherence for this sub-population. Sex hormones play integral roles in appetite regulation [31, 30]. Although we did not analyze hormones directly, it is possible that these signals might be differentially stronger than other anorexigenic signals originating from the gut for women compared to men, possibly explaining some of the sexual dimorphism in our results. Additionally, self-reporting biases might have influenced the 24-hr dietary recalls, and thus our dietary compliance measures, differentially for men and women [29]. We accounted for this by calculating macronutrient intake as a percentage of total energy intake, but these biases were not assessed directly in the survey instrument and thus we were unable to specifically adjust for them.

Plasticity of the gut microbiota over the first ten weeks of the dietary intervention was higher for VS compared to US subjects on either diet, but significant only for low-carb dieters. Again, we saw strong diet- and sex-specific differences between plasticity and dietary compliance. Although higher microbiota plasticity was seen in VS low-carb dieters, there was no correlation between plasticity and dietary compliance for either sex, implying that for diets with restricted carbs the microbiota might play a larger role in the proportion of nutrients extracted from foods and less of a role in appetite suppression. The correlations between 10-week plasticity and dietary compliance were positive for men on the low-fat diet and negative for women, again implying that sex-specific differences are likely in our measurement of dietary compliance and/or in the physiological impacts of gut microbiota on appetite suppression.

Plasticity of the gut microbiota over ten weeks, and in response to the intervention, was higher for both diets than daily plasticity at either time point. This was expected as dietary changes can have strong influences on taxonomic composition and functional capability of the gut microbial community [21, 19, 38, 56, 45]. However, the non-standardized, self-titrated interventions did not lead to a convergence of subjects to similar gut microbiota communities. After 10 weeks on the dietary intervention, subjects’ community membership and structure were still more similar to their own pre-diet communities rather than to other subjects on the same diet. The repeated measures at both time points (pre-diet and 10 weeks) enabled us to observe that the plasticity of the microbial community in response to the diet was higher than daily variability at either time point. As we were unable to evaluate the robustness of these plasticity findings in our validation cohort due to differences in sampling regimes, there remains the need for other studies to corroborate these plasticity findings. Future analyses should be stratified by sex, with specific attention paid to menopausal state of women included.

Finally, our study highlighted the importance of validation cohorts in microbiota studies with small to moderate sample sizes. Contrary to our hypothesis, we found no specific taxonomy-related microbial features clearly associated with 12-month weight-loss success. This could be due to our small sample size (n=25-34 for any diet-cohort combination), indeed, only ~ 20% of ASV-clusters were detected in at least half of the subjects on either diet, limiting our statistical power to identify differences between VS and US subjects. Although our sample size was comparable to other published studies investigating the gut microbiota and diet responsiveness [32, 38, 18], our analysis included stringent filtering of samples with at least 10,000 reads and ASV-clusters present in at least 10% of subjects on the focal diet. We also included replicate samples for most subjects in order to reduce false positive identification during differential abundance testing. The majority of pre-diet ASVs and ASV-clusters identified as differentially abundant between weight-loss groups on the low-carb diet were discordant across the discovery and validation cohorts in both magnitude and direction. In addition, our finding from the discovery cohort that subjects on the low-carb diet with higher pre-diet P/B ratio had higher 12-month weight loss was not confirmed in the validation cohort. These inconsistencies could be the result of the significantly lower dietary compliance and weight loss achieved by low-carb dieters in the validation cohort compared to subjects from the discovery cohort. The inclusion of a validation cohort illustrated the challenges in generalizing findings from an initial modest sample size to even a similar population. Future research, specifically designed and powered for the outcome of interest, will be necessary to determine if specific taxa or higher P/B ratio are important for 12-month weight loss in larger populations.

## Conclusions

Our study is the first to examine the connection between gut microbiota and 12-month weight loss success on two common diets – low-carb and low-fat – and on the intermediary of dietary compliance. For subjects on the low-fat diet, higher weight loss was observed in subjects with higher pre-diet daily microbiota plasticity but this appears to be mediated by adherence only for men. Low-carb dieters appear to obtain little benefit from daily pre-diet gut microbiota plasticity in terms of weight loss. Higher plasticity of gut microbiota over ten weeks, and in response to the dietary intervention, was seen for subjects who lost the most weight after 12 months of low-carb dieting, but we found no correlations for this group between plasticity and dietary compliance. Despite strong correlations between gut microbiota plasticity over ten weeks and dietary compliance for low-fat dieters, no significant difference in plasticity was noted between VS and US weight loss success groups.

Gut microbiota plasticity has not been extensively studied in relation to sustained weight loss or dietary adherence. Here we present data that suggests the plasticity of the gut microbiota may be related to both in a sex-and diet-specific manner. Our work highlights the importance of investigating kinetic trends in gut bacterial community composition, including long-term shifts and daily plasticity, in addition to explicitly seeking static microbiota signatures and patterns, when studying individual differences and predisposition to successful weight loss.

## Methods

### Study population and sample collection

Overweight and obese adults in the Diet Intervention Examining The Factors Interacting with Treatment Success (DIETFITS) study enrolled between Fall 2013 and Spring 2014 were approached for inclusion in this study. DIETFITS was a 12-month randomized clinical trial of low-carb and low-fat diets [57, 24] (clinical trial registration NCT01826591). The diets had no specific caloric, fat or carbohydrate restrictions, but instead involved counseling sessions focused on three main components central to sustaining a low-carb or low-fat diet. First, for the initial eight weeks, subjects were instructed to progressively reduce either carbohydrate or fat intake as much as possible (with an objective of achieving 20 g/day), and maintain their lowest possible intake for at least several weeks. Second, after the initial eight weeks they were then encouraged to titrate their intake by increasing fat or carbohydrate consumption by increments of 5-15 g/day and maintaining that intake for a week or more while noting their satisfaction (e.g., satiety, palatability) and weight loss success. After settling on a sustainable target (one that left them feeling full but still able to lose weight), they were asked to maintain that intake level for the remainder of the study. The third component of the study’s strategy was promotion of a high-quality diet focused on whole, real foods, that are mostly prepared at home and that contain as many vegetables as possible. Subjects were assigned to attend 22 in-person instructional sessions related to nutrition, behavior, emotions, and physical activity. Participant data, including clinical outcomes, was collected and managed using REDCap electronic data capture tools hosted at Stanford University [27]. Subjects from the discovery cohort were asked to provide self-collected fecal samples from three consecutive days at two separate time points (pre-diet and 10 weeks after initiation of the dietary intervention). Subjects from the validation cohort were asked to provide two self-collected pre-diet fecal samples. Fecal samples were stored at -20°C until delivered to the lab and then at -80°C until processing. A total of 424 fecal samples with sufficient sequencing depth in high-quality reads were collected from 66 discovery cohort and 56 validation cohort subjects who provided complete weight data.

### DNA extraction and 16S rRNA gene sequencing

DNA was extracted from 50-150mg fecal material using the Qiagen PowerSoil DNA Isolation Kit (Qiagen, Venlo, The Netherlands) with the following protocol modifications: samples were incubated in lysis buffer at 65°C for 10 minutes, bead beating was conducted for 20 minutes, and all subsequent vortexing steps were replaced with gentle but thorough inversions. Extracted DNA was stored at -20°C. The V4 region of the 16S rRNA gene was amplified using the protocol of Caporaso et al. [13]. Briefly, samples were amplified in triplicate 25ul PCRs in 96-well plates with the final volumes per reaction: water 10.9 ul, MasterMix (5 PRIME HotMasterMix) 10 ul, reverse primer 0.1 ul, forward primer 1 ul, template DNA 3 ul. Replicates were run simultaneously on 3 thermal cyclers for 30 cycles of: 94°C 45s, 52°C 60s, 72°C 90s with a ten minute extension at 72°C at the end. Amplification was verified by gel electrophoresis on pooled replicates (failures were repeated) and bands were excised and cleaned (MO BIO UltraClean-htp 96 Well PCR Clean-Up Kit) per manufacturer protocol. DNA was quantified (Invitrogen Quant-iT ds DNA Assay Kit, High Sensitivity) using a microplate reader (FLEXstation II 384-Fluorescent 6), and amplicons were combined in equimolar ratios. This pooled DNA was concentrated via ethanol precipitation and then resuspended in nuclease-free water. DNA libraries were submitted to the Functional Genomics Facility at Stanford University for sequencing on an Illumina MiSeq (2 x 250bp) over 7 separate runs, yielding 73.3M raw reads.

### DADA2 Amplicon Sequence Variant (ASV) Sample Inference and Tree Building

The `DADA2` R package [10] was used for quality filtering, denoising, chimera removal and sequence inference (obtaining sequence counts). Forward reads were trimmed to length 240 in all but one sequencing run (length of 230 due to lower quality); reversed reads, commonly with lower quality, were trimmed to length ranging from 160-200 across sequencing runs. After merging, default settings were used for error estimation and denoising with the following exception: `maxEE` = 2, meaning that merged reads with expected error higher than `maxEE` were discarded (where *EE* = Σ10^−*Q*/10^). Finally, after amplicon count inference, chimeras were removed and sequences of length 230-234 bp were retained; 37.5M (51%) reads passed filtering criteria. Taxonomic assignment of the 4,234 unique sequences was performed according to the Bioconductor workflow [10, 9] using RDP trainset 16 [63] and Silva v128 [51] databases.

#### Phylogenetic tree estimation

In accordance with the Bioconductor workflow [9], two R packages were used to estimate a phylogenetic tree from obtained sequences. `DECIPHER` [66] was used to perform multiple alignment, and `phangorn` [54] to fit a Generalized time-reversible Gamma rate variation (GTR+G+I) maximum likelihood tree initialized at the neighbor-joining tree (parameters *k*=4 and inv=0.2 were used for `phangorn::pml` function).

#### Sequence clustering using phylogeny

Due to high resolution of the `DADA2` pipeline, the obtained amplicon sequence variant (ASV) count-matrix was very sparse. In order to attain more overlap in organism counts between samples and subjects, we clustered ASVs into phylogenetically similar sequence groups using the estimated phylogenetic tree. These clusters were generated by cutting an associated phylogenetic tree at the height, *h* = 0.1, corresponding to median difference between member sequences equal to 7.5 bp (out of 233 bp). This is equivalent to 96.8% sequence identity, which correlates with thresholds to differentiate genera [5, 61]. We used both the original ASV data and ASV-clusters for downstream analysis. This ASV-clustering approach is more accurate than the traditional OTU clustering, as the phylogenetic distance is considered in the sequence grouping process.

### Statistical Analyses

#### Ordination

To visualize the data we used Principal Coordinate Analysis (PCoA) with Bray-Curtis distance applied to inverse-hyperbolic-sine (asinh) transformed count data. The transformation prevents the Bray-Curtis dissimilarity metric from placing too much weight on species/ASVs highly abundant in all samples. All samples were included together in the computation of the ordination projection. The plots were then faceted into distinct dietary intervention assignments. Differences in community composition were tested using PERMANOVA (`adonis` function from the `vegan` package [48], permutation = 999).

#### Microbiota Diversity and Plasticity

Gut microbiota phylogenetic *α*-diversity was estimated using the procedure of Nippress et al. [47], based on rarefaction curves. Phylogenetic diversity was evaluated at a depth of 11,000 sequence reads – the level of the minimum library size of samples post-filtering.

Daily pre-diet gut microbiota variability was estimated using pairwise *β*-diversity between each subject’s pre-diet samples. If a subject provided three samples, only the pairwise *β*-diversity for consecutive days were used to compute the average. Consecutive samples in the discovery cohort were taken a median of 1 day apart (IQR 1, 2) with an exception of a few subjects who provided samples up to 5 days apart. The time span between an individual subject’s samples was balanced across weight-loss groups. To estimate variability between gut microbiota pre-diet and 10 weeks into the dietary intervention, we calculated the pairwise *β*-diversity between each pre-diet and 10-week sample for each subject, and then calculated the mean across all pairwise comparisons. For all plasticity measures, Jaccard dissimilarity and unweighted UniFrac distance were used to asses the variability in taxa presence/absence, whereas Bray-Curtis dissimilarity, weighted UniFrac distance, and Jensen-Shannon divergence were used to estimate variability in abundances. The difference in variability between VS and US subjects was tested for significance separately for each diet using non-parametric Wilcoxon rank-sum test.

#### Differential abundance testing

We looked for pre-diet ASV-clusters, and single ASVs, that were differentially abundant with respect to 12-month weight loss. The inclusion of consecutive daily samples (repeated measures) for subjects allowed for a more accurate estimation of the underlying processes and helped reduce false discoveries. The `limma` differential abundance (DA) testing framework [52] is the most conservative analysis that allowed us to model variability of repeated measurements as within-subject random effects. Before using `limma` we transformed the raw count data by first computing the library size factors using `estimateSizeFactors` from `DESeq2` package [43] with the argument `type “poscounts”`, which was specifically developed for sparse sequencing data. We used a customized version of `limma::voom` function, where we substituted the log2-counts per million (logCPM) transformation intended for bulk human RNA-seq data, with an inverse-hyperbolic-sine 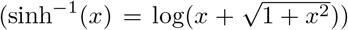 transformation shown to be more appropriate for data which follows a negative binomial distribution (as is the case for 16S rRNA gene sequencing) [40, 34, 7, 35]. Functions for differential abundance estimation from `limma` were then used to fit the pre-diet ASV or ASV-cluster abundances. For increased modeling accuracy and higher power, the method was applied to all samples from both diets together. Testing was performed by evaluating the contrast between weight-loss success categories within each diet separately (using diet-weight loss interaction terms in the model design). An additional fixed-effect term corresponding to sample sequencing “lane” assignment was included in the model to account for batch effects from different sequencing runs. For each of the two diets we tested both the contrast between subjects from the VS compared to US weight-loss group, and also conducted secondary analyses modeling the outcome as a continuous variable (percent weight lost at 12-months). Only the taxa found in at least 10% of subjects on the given diet and significant after adjusting for I multiple hypothesis testing (Benjamini-Hochberg method [4]) at level *α* = 0.05 were retained.

## Declarations

### Acknowledgements

We are grateful to Daniel Sprockett and Elizabeth Costello for their critical comments on this manuscript. We would like to thank the Stanford Research Computing Center for providing computational resources (Sherlock cluster) that contributed to these research results. We thank the large team of researchers who executed the DIETFITS study, specifically Jennifer Robinson, Lisa Offringa, Erin Avery, Joseph Rigdon, and Lucia Aronica, for their assistance in obtaining and interpreting participant data. We also acknowledge the study participants without whom this investigation would not have been possible.

### Funding

This study was supported by a grant 1R01DK091831 from the National Institute of Diabetes and Digestive and Kidney Diseases, funding from the Nutrition Science Initiative, and a gift from Robert and Mary Ellen Waggoner. JAG was funded by a Stanford Interdisciplinary Graduate Fellowship. LHN was funded by NIAID R01 AI112401 to SPH. Research IT grant support was provided by Stanford Clinical Translational Science Award number UL1 TR001085 from NIH/NCRR.

### Availability of data and materials

R scripts used for the analysis are available in Additional Files 1-6. All sequencing reads will be deposited in the Sequence Read Archive under BioProject number SUB5292808.

### Author’s contributions

JP and CDG conceived of and designed the project. TDH conducted all wet lab work. SH provided conceptual advice on analysis of data. JAG and LHN analyzed and interpreted the data. JP, CDG, SPH supervised research. JG prepared the manuscript, with contributions from all authors. All authors read and approved the final manuscript.

### Ethics approval and consent to participate

This research was approved by the Panel on Human Subjects in Medical Research (protocol 22035) at Stanford University, Stanford, CA, USA. All participants provided written informed consent.

### Competing interests

The authors declare that they have no competing interests.

---

**Figure S1:**
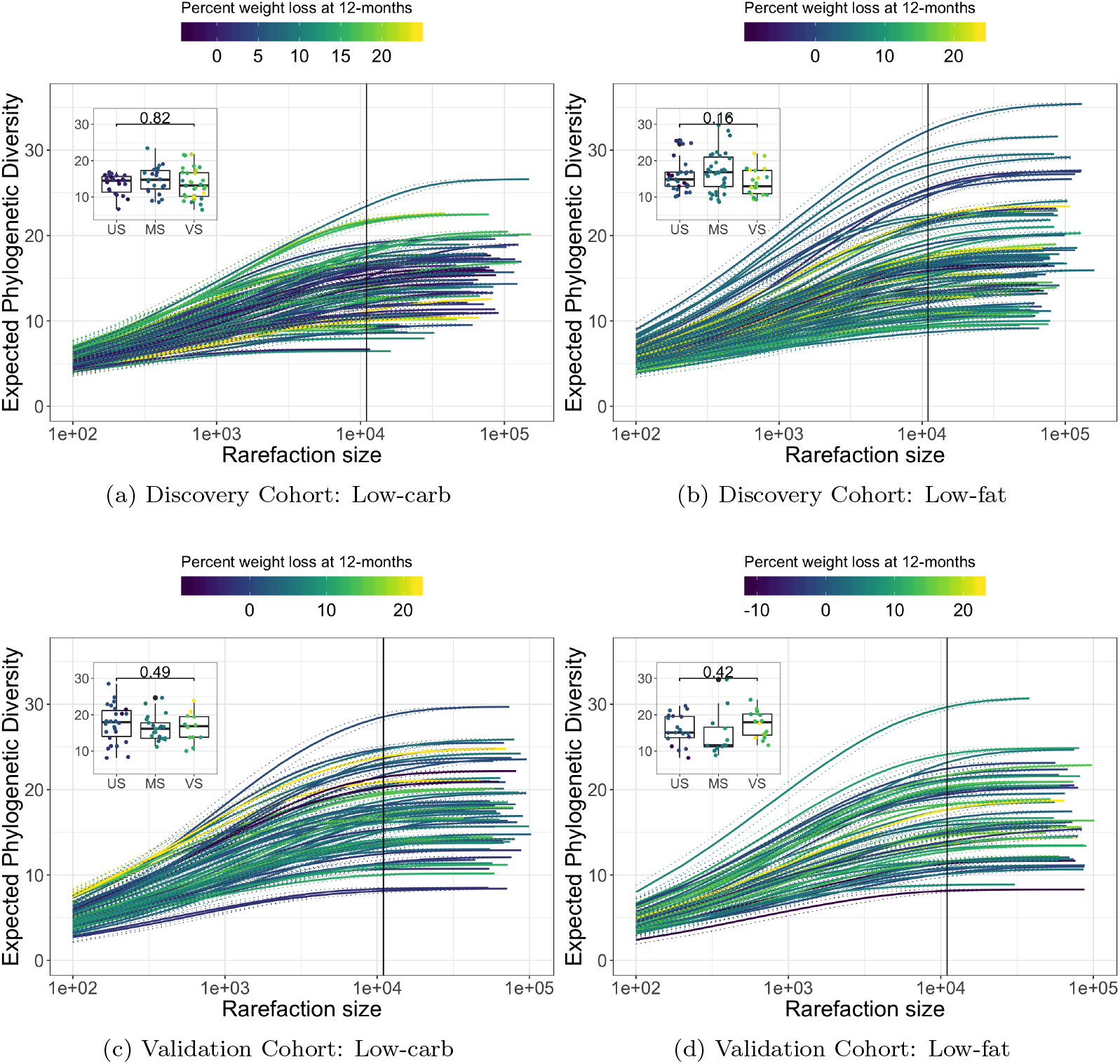
Phylogenetic *α*-diversity in pre-diet samples. Rarefaction curves for low-carb (*left*) and low-fat (*right*) diet, separated by discovery (*top*) and validation (*bottom*) cohorts. Curves were computed using the methods described in [47]. Inset boxplots report non-significant differences in *α*-diversity (at rarefaction level = 11,000 reads) between subjects from the VS and US 12-month weight loss success categories.

**Figure S2:**
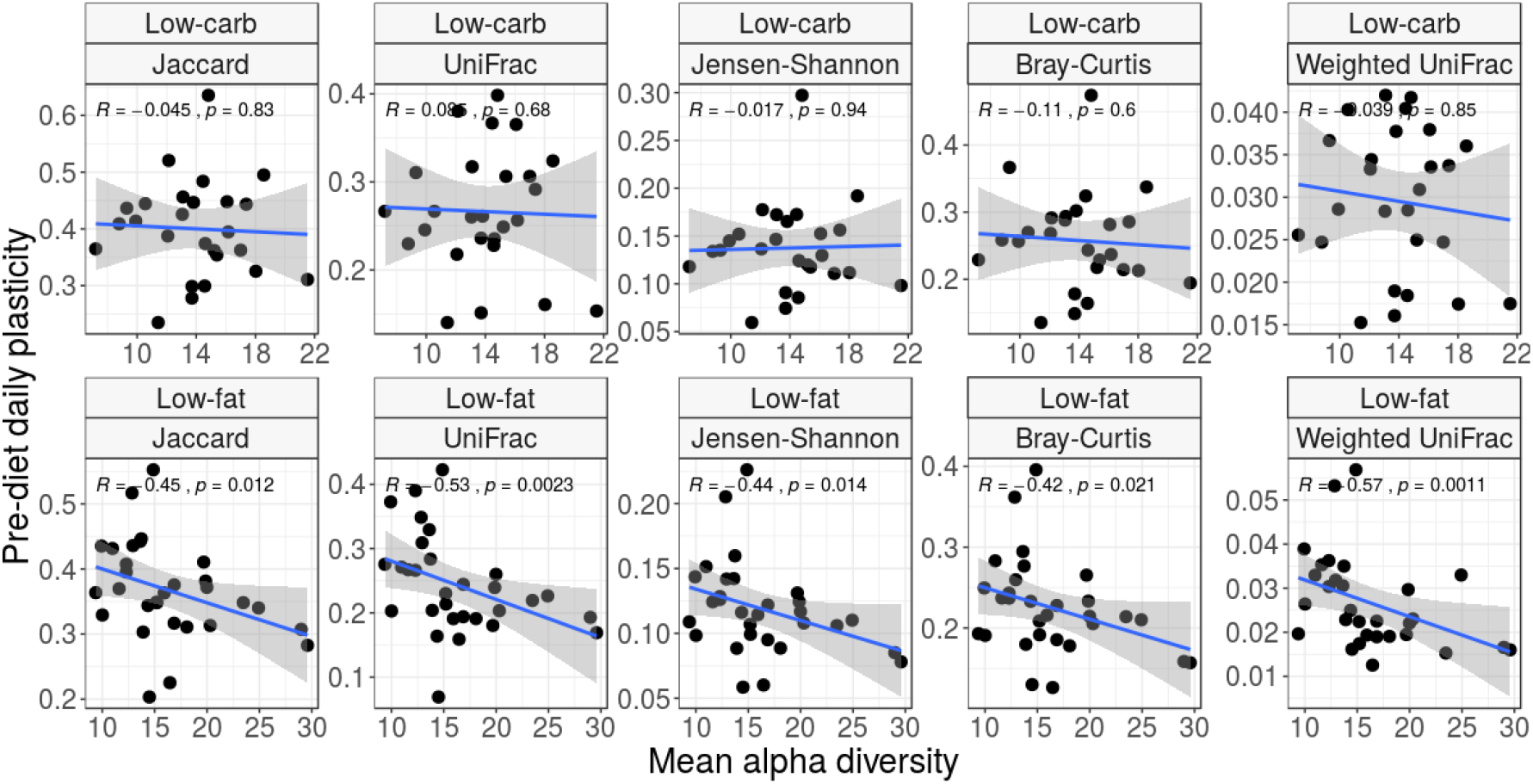
Beta diversity negatively correlated with alpha diversity. Mean baseline bacterial community diversity levels are negatively correlated with baseline bacterial plasticity for subjects on Low-fat diet.

**Figure S3:**
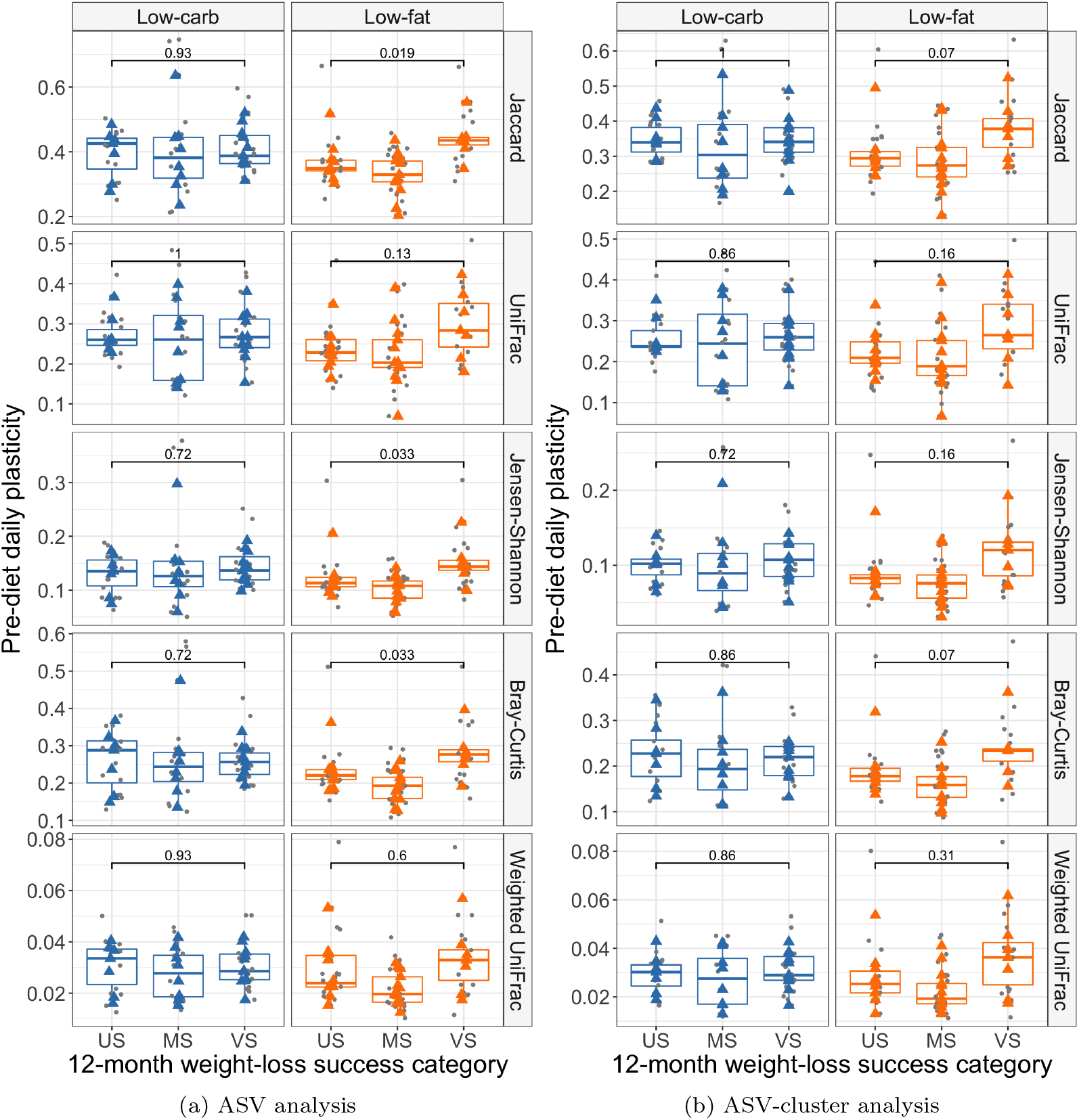
Day-to-day plasticity (*β*-diversity) between pre-diet fecal samples from subjects in the discovery cohort, grouped by 12-month weight loss success. Jaccard, unweighted UniFrac, Jensen-Shannon-Divergence, Bray-Curtis, and weighted UniFrac distances are shown for low-carb diet (left panels, in blue) and low-fat diet (right panels, in orange). Grey points indicate computed pairwise dissimilarities between samples; colored (low-carb and low-fat) points indicate the average sample-to-sample dissimilarity for each subject.US - Unsuccessful, < 3% weight loss; MS – Moderately successful, 3 – 10% weight loss; VS – Very successful, > 10% weight loss. P-values shown for Wilcoxon rank-sum test comparing US and VS groups. The analysis was done at the level of individual ASVs (a) and also ASV-clusters (b).

**Figure S4:**
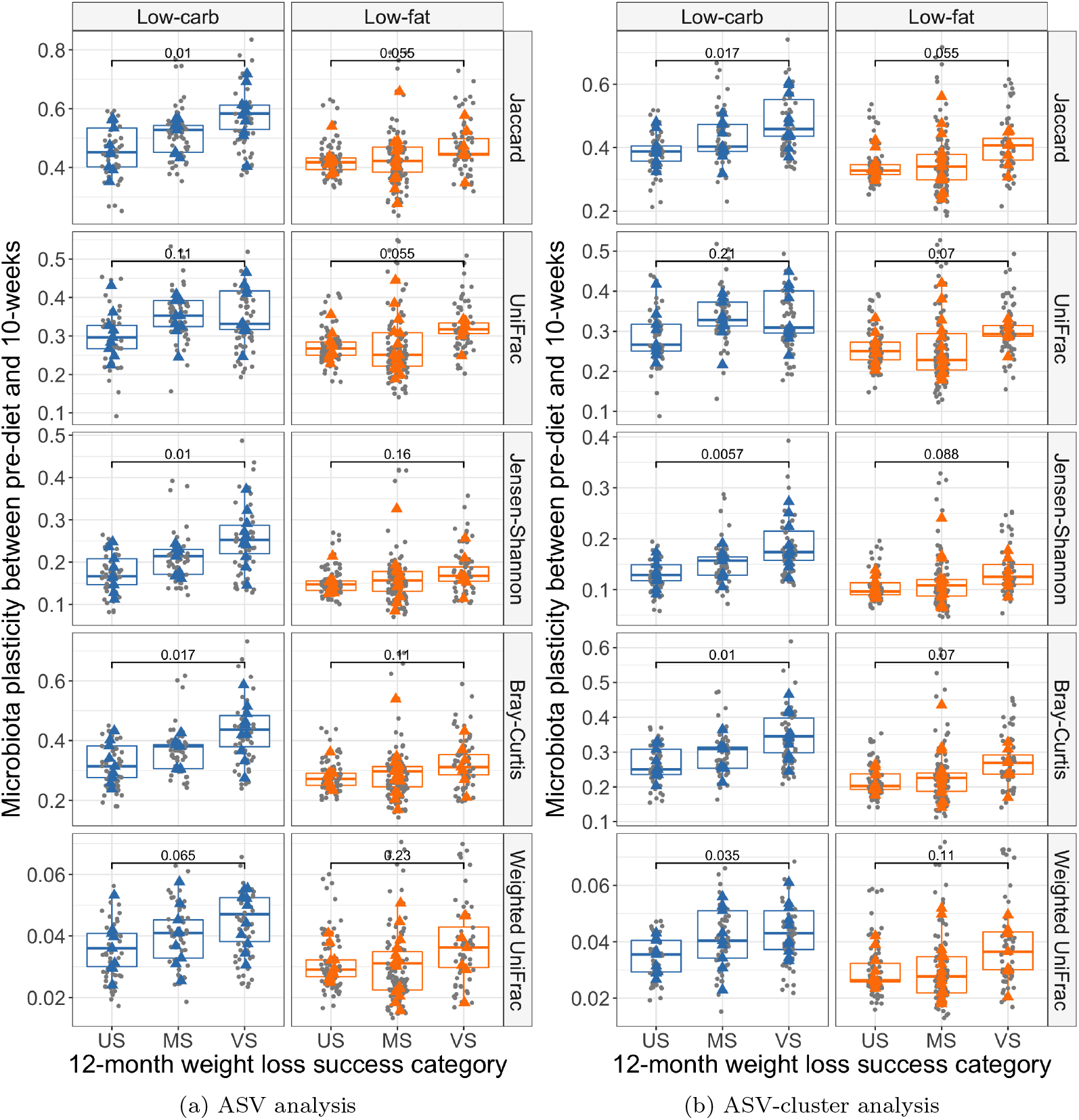
Microbial community composition shift over ten weeks, and in response to dietary intervention. *β*-diversity between pre-diet (baseline) and post-diet samples (taken at 10 weeks after initiation of the dietary intervention) from each subject was computed using Jaccard, unweighted UniFrac, Jensen-Shannon-Divergence, Bray-Curtis, and weighted UniFrac distances. Results shown for discovery cohort subjects on a low-carb (left panels, in blue) or low-fat (right panels, in orange) diet. Grey points indicate each computed pairwise dissimilarity between samples; colored points correspond to the average baseline-to-10-week plasticity for each subject. US – Unsuccessful, < 3% weight loss; MS – Moderately successful, 3–10% weight loss; VS – Very successful, ≥ 10% weight loss. P-values shown for Wilcoxon rank-sum test comparing US and VS groups. The analysis was done at the level of individual ASVs (a) and also ASV-clusters (b).

**Figure S5:**
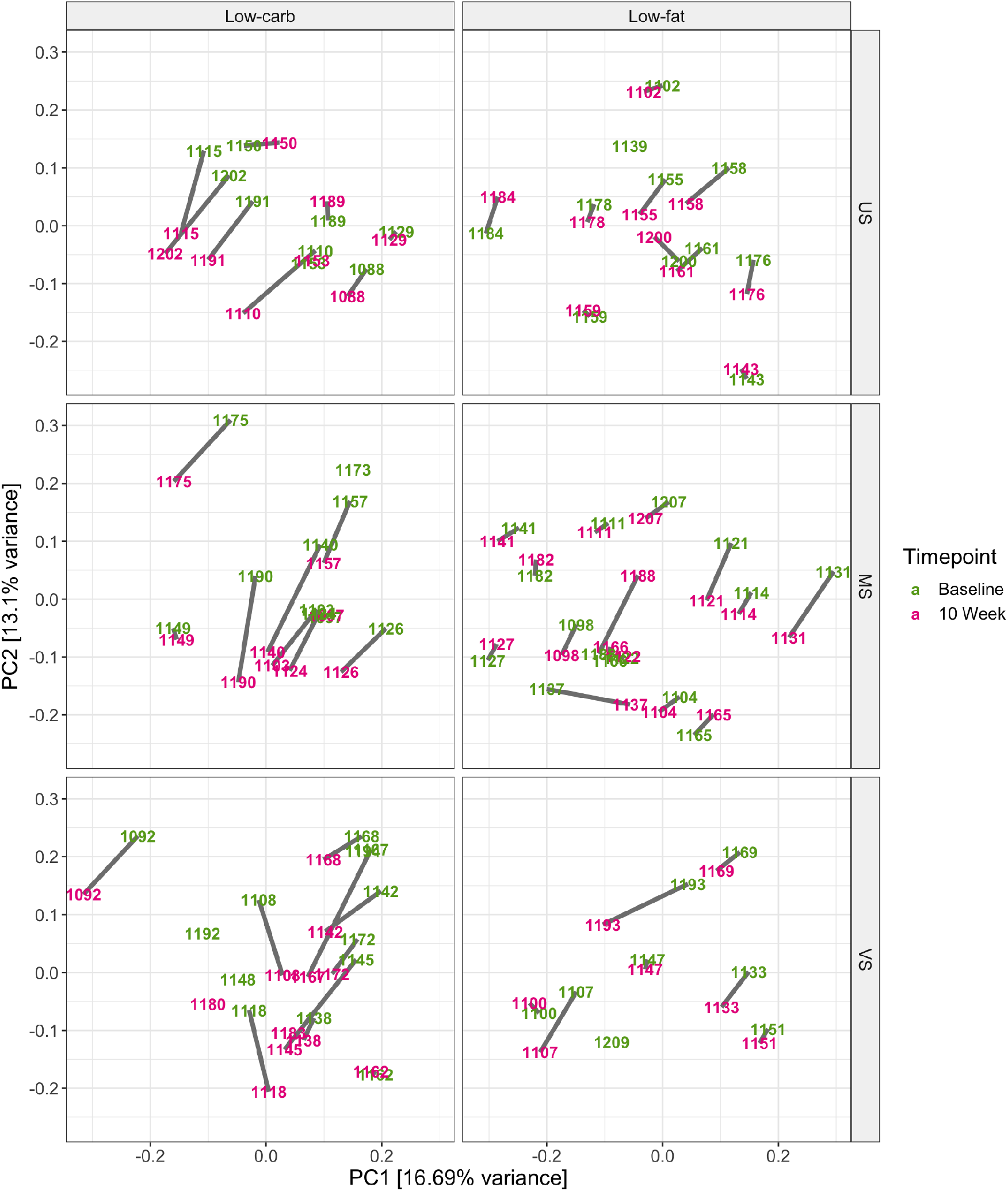
Microbiome shift in response to dietary intervention. Subject centroids (coordinates averaged over sample replicates) before (green) and 10 weeks after the start of the intervention (purple) for low-carb (*left*) and low-fat (*right*) diet. Data points are labeled with corresponding unique subject ID. Vertical panels correspond to weight loss success categories – very successful (VS), moderately successful (MS) and unsuccessful (US).

**Figure S6:**
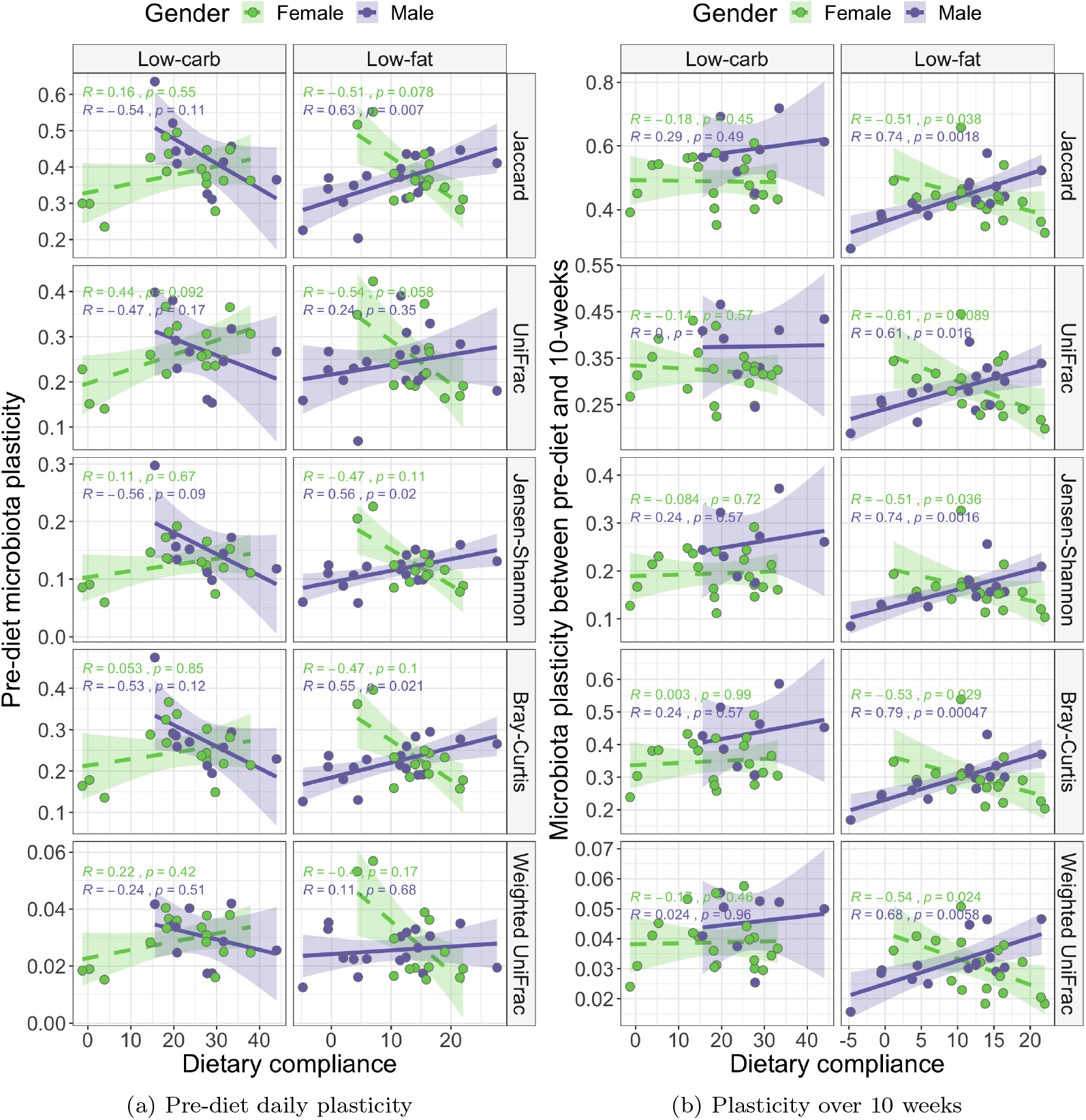
Sex- and diet-specific correlation between microbial community composition plasticity and dietary compliance. Plasticity across various *β*-diversity measures between daily pre-diet samples (a) and between pre-diet and 10-week samples (b) from each subject compared to dietary compliance. Results shown for discovery cohort subjects on a low-carb (left panels) or low-fat (right panels) diet. Male (purple) and female (green) subjects show opposite correlations in many cases. Spearman’s rank correlation coefficients are shown for each subgroup.

**Figure S7:**
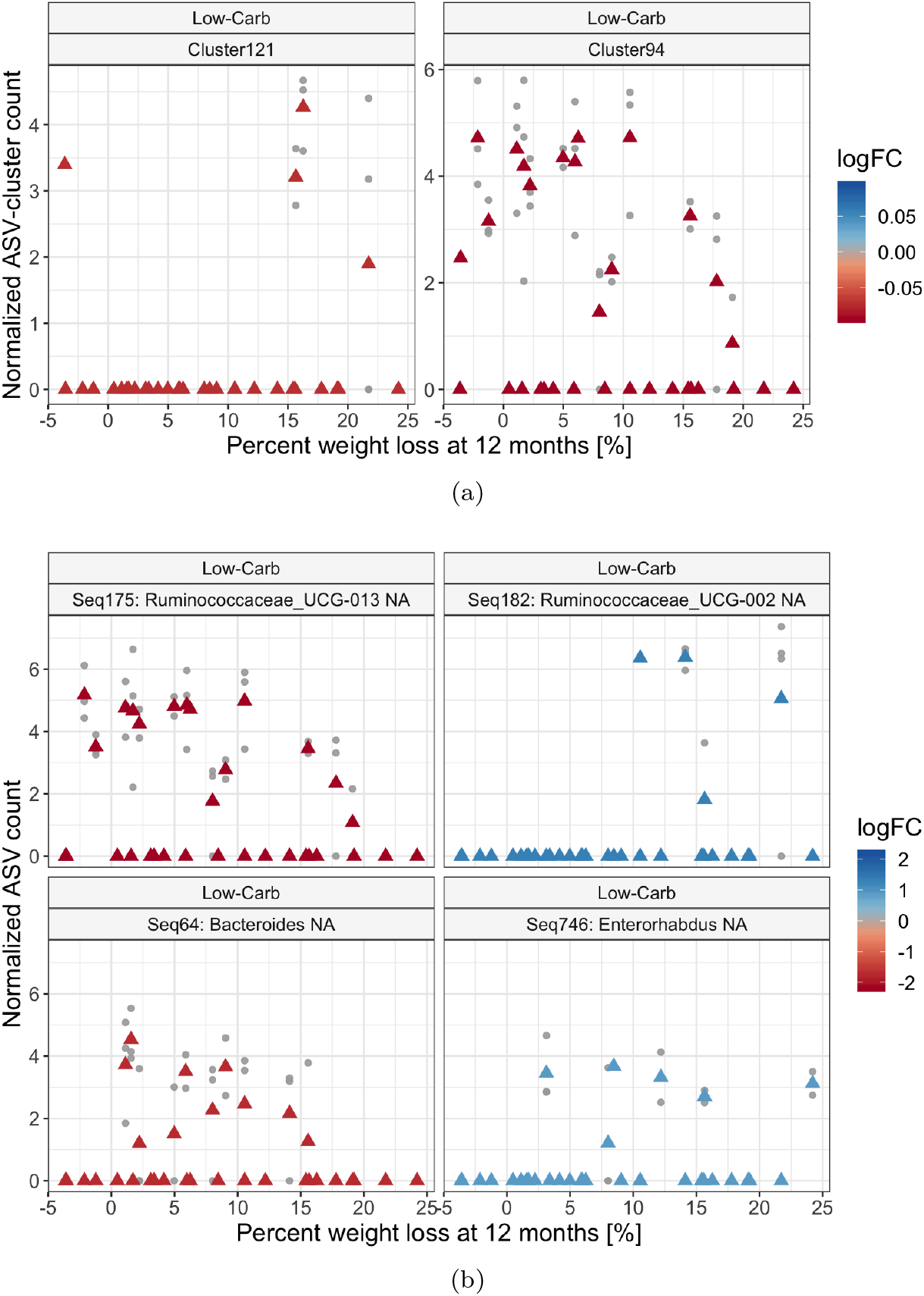
Taxa identified differentially abundant for subjects on low-carb diet. Analyses were conducted with ASV-clusters testing for difference in abundance across a continuous response variable – percent weight loss (a) and also with individual ASVs and tested for contrast between categorical weight-loss groups: VS compared to US subjects. Note: Cluster94 contains Seq175, as can be seen in the similarity of plots. Individual ASVs and ASV-clusters were normalized and asinh-transformed for variance stabilization prior to analysis; the normalized, transformed values are shown on the y-axis. Grey points represent individual samples and triangles represent the mean value for each subject. No taxa were found differentially abundant on the low-fat diet for either analysis.

